# Gene function is a driver of signaling pathway evolution following whole genome duplication

**DOI:** 10.1101/2023.02.03.527009

**Authors:** Jasmine A. Richman, Leah R. Davis, Michael P. Phelps

## Abstract

The genome of many plant and animal species are heavily influenced by ancestral whole genome duplication (WGD) events. These events transform the regulation and function of gene networks, yet the evolutionary forces at work on duplicated genomes are not fully understood. Genes involved in cell surface signaling pathways are commonly retained following WGD. To understand the mechanisms driving functional evolution of duplicated cell signaling pathways, we performed the activin receptor signaling pathway in rainbow trout (RBT). Rainbow trout are a model salmonid species that exhibit a duplicated genome as a result of an ancestral WGD that occurred in all teleost fish, and a second more recent WGD found in salmonid fishes. This makes RBT a powerful system for studying ohnolog evolution in a single species. We observed that regulation of the duplicated activin receptor signaling pathway is commonly driven by tissue-specific expression of inhibitors and ligands along with the subfunctionalization of ligand ohnologs. Evidence suggests that for inhibitors and R-Smad signaling molecules, there is ongoing pressure to establish a single copy state which may be driven, in part, by regulatory suppression of select ohnologs. The core transmembrane receptors and Co-Smad signaling cascade members are high duplicated yet exhibit contrasting expression dynamics where receptors tend to share expression across tissues while dominance of a single ohnolog is common for the Smad4, Co-Smad gene family. Our findings provide support for a generalized model where gene function and gene dosage have a complementary role in ohnolog evolution following WGD.

## Introduction

Whole genome duplication events (WGD) are hypothesized to play a role in driving speciation by providing a template for the diversification of gene function. While most duplicated genes are eventually lost following WGD, specific classes of gene families are more likely to retain duplicated members (Lien et al. 2016; Birchler and Yang 2022). This includes genes involved in intercellular signaling, gene transcription, and the formation of core protein complexes (Huminiecki et al. 2009; Huminiecki and Heldin 2010; Lien et al. 2016).

Understanding the forces driving the expansion of gene families involved in these important cellular processes is critical to defining the role WGDs play in the diversification of multi-cellular life.

Two ancestral WGD events have influenced the construction of the vertebrate genome (Huminiecki and Heldin 2010). In tetrapods, further duplication of the genome is generally not tolerated, possibly due to the associated genomic instability resulting from a WGD. This is not the case for teleost fish, which have experienced additional genome duplication events throughout their lineage and can endure further genome expansion either through natural processes or experimental manipulation (Pandian and Koteeswaran 1998). Teleost fish are the most diverse vertebrate lineage, with over 30,000 species, and their WGD events are hypothesized to have contributed to their rapid diversification (Volff 2005; Santini et al. 2009). In addition to the teleost specific WGD (Ts3R) that occurred approximately 320 million years ago, salmonids experienced another autotetraploidization WGD event (Ss4R) approximately 80-100 million years ago (Lien et al. 2016). The presence of two WGD events in salmonids, one ancestral and one more recent, have establish salmonid fishes as a unique model for investigating the evolution of gene ohnologs after WGD (Berthelot et al. 2014; Lien et al. 2016; Gundappa et al. 2022).

Following WGD, changes in the coding sequence, expression level, and tissue distribution of ohnologs can occur to maintain or develop novel functions necessary for adaptation or the evolution of new traits. This functionalization process can cause the degeneration of gene family ohnologs back to a single gene state (defunctionalization), allow them to establish partial functional redundancy (hypofunctionalization or subfunctionalization), or to adopt completely new functions (neofunctionalization; Force et al. 1999; Birchler and Yang 2022). While the drivers of these evolutionary outcomes are not fully understood, current theory suggests that one such pressure is the need to balance stoichiometric levels of gene products that are involved in dosage sensitive cellular processes (Gillard et al. 2021; Birchler and Yang 2022). Intercellular signaling pathways are dosage sensitive and have essential functions in the growth and development of tissues, which influence organismal physiology and body plan organization (Huminiecki et al. 2009; Harper et al. 2021). Since cellular communication is dependent on strict control of the timing and level of signaling pathway components, rediploidization of cell signaling gene ohnologs presents a fundamental challenge to maintaining proper intercellular communication. This is due to the fact that functional changes in one signaling pathway member influences the evolutionary outcome of other members in the pathway (Freeling and Thomas 2006; Huminiecki and Heldin 2010; Robertson et al. 2017).

A classic example of a signaling pathway that has evolved increased complexity with successive WGD events is the transforming growth factor beta (TGF-β) superfamily. The TGF- β superfamily arose in early metazoans and is characterized by significant expansion throughout evolutionary time as new members, added through genome duplication events (1R and 2R), adopt unique functions (Huminiecki et al. 2009). The superfamily consists of four major pathways; nodal, TGF-β, bone morphogenetic protein (BMP), and the activin signaling pathway, each characterized by the type of receptors that mediate signaling. Similar to other pathways within the broader TGF-β superfamily, the activin receptor signaling pathway functions to regulate the growth and differentiation of numerous cell types. The pathway signals primarily through autocrine or paracrine mechanisms functioning within specific tissues. Dimeric TGF-β superfamily ligands bind to a tetrameric complex of type I and type II serine-threonine kinase receptors. Ligand binding activates a serial phosphorylation cascade whereby type II receptors phosphorylate type I receptors, which in turn phosphorylate receptor activated Smad signal transduction proteins (R-Smads), which translocate to the nucleus to modify gene transcription. Activation of the signaling pathway is controlled in part by inhibitory molecules that interfere with ligand binding, receptor activation, or signal transduction. The precise combination of ligands, receptors, and inhibitors determines the response of cells to activin receptor signals and therefore it is the interaction and level of each pathway member that influences the overall function of the pathway (Massagué 2012). Disruption of this balance has been shown to have significant consequences for vertebrate physiology (Lee et al. 2005; Pangas et al. 2007; Phelps et al. 2013).

The activin receptor signaling pathway is essential for embryonic and reproductive development in mice and is arguably the most important regulator of muscle fiber size in mammals (McPherron and Lee 1997; Lee and McPherron 2001; Lee et al. 2012). While less is known of the functions of the activin receptor signaling pathway in teleost fish, its function in muscle growth appears to be conserved (Medeiros et al. 2009; Phelps et al. 2013; Khalil et al. 2017; Kishimoto et al. 2018). As a result of the two salmonid WGD events (Ts3R and Ss4R), all of the activin receptor signaling pathway genes remain duplicated in rainbow trout (RBT, *Oncorhynchus mykiss*), a model salmonid species. In fact, just under half (42%) of RBT activin receptor signaling pathway ohnologs have retained all four duplicated forms. It currently is unclear how duplication of the activin receptor signaling pathway has influenced the organismal biology of salmonids compared to other fish and tetrapods.

Traditional methods of investigating the evolution of gene ohnologs often analyze genes across species, within individual gene families, or genome wide. Examining gene evolution across species can be influenced by the specific selective pressures or life histories exhibited by each species, which may mask fine scale drivers of ohnolog evolution. Similarly, focusing on specific families of genes or broad genome trends in ohnolog evolution while failing to account for the interactive gene networks by which those genes function can miss critical information about how cellular processes influence ohnolog evolution. To gain insight into this important evolutionary process, we capitalized on the unique evolutionary history of salmonid fishes to characterize the evolutionary mechanisms driving intercellular signaling pathway evolution after multiple WGD events in a single species, across an entire interconnected biological pathway.

This was accomplished by developing a detailed, quantitative gene expression atlas of 49 activin receptor signaling pathway ohnologs across 23 adult RBT tissues. Combining the level and tissue specific expression patterns of these genes with molecular evolutionary analysis of activin pathway gene ohnologs provided insight into the role gene duplications play in driving functional divergence of the activin receptor signaling pathway across both the Ts3R and Ss4R WGDs in salmonids. Our findings provide support for the significant role gene function plays in driving the evolution of ohnologs after WGD and that, while important, gene dosage alone cannot fully explain the processes underlying ohnolog retention or loss.

## Results and Discussion

### Duplication of the Activin Receptor Signaling Pathway in Rainbow Trout

To gain insight into how protein function influences the evolution of interconnected gene products after WGD, we examined the functional evolution of the activin receptor signaling pathway in RBT (*O. mykiss*) across the ancestral teleost Ts3R and salmonid lineage specific Ss4R genome duplication events found in this species. The activin receptor signaling pathway is a member of the TGF-β superfamily which has critical roles in organismal growth and development. While a reference genome exists for RBT, most ohnologs have not been identified or named according to their evolutionary relationships. We therefore used molecular phylogeny to identify and name 59 genes involved in the activin receptor signaling pathway (Table 1). The characterized genes represented 19 distinct gene families with proposed functions as: serine-threonine kinase receptors, ligands, signal transducers, and inhibitors. It is important to note that some pathway members, such as activin C (*inhbc*) and *bambi*, have poorly defined functions and may have both stimulatory and inhibitory functions depending on the tissue context, but are primarily regarded as pathway inhibitors (Gold et al., 2009; Onichtchouk et al., 1999). The proposed nomenclature used in naming the various pathway members followed established naming conventions that refer to Ts3R ohnologs alphabetically (e.g., a and b), based off their ancestral chromosome origins and sequence similarity to the northern pike (*Esox lucius*), and Ss4R ohnologs numerically (e.g., 1 and 2) using their similarity to the Ts3R ohnologs (Table 1; Parey et al. 2022).

**Table 1.** Proposed Nomenclature of Activin Receptor Signaling Pathway Ohnologs.

| Gene Name | Gene ID | Protein Accession | Nucleotide Accession | Chromosome | Ancestral Chr. | Expression Probe |
| --- | --- | --- | --- | --- | --- | --- |
| acvr1aa1 | LOC110501518 | XP_021434832 | XM_021579157 | 22 | 3a | Yes |
| acvr1aa2 | LOC110520052 | XP_021452705 | XM_021597030 | 3 | 3a | Yes |
| acvr1ab1 | LOC110495748 | XP_021426823 | XM_021571148 | 18 | 13b | Yes |
| acvr1ab2 | LOC110528876 | XP_021466722 | XM_021611047 | 7 | 13b | Yes |
| acvr1ba1 | LOC110532699 | XP_021472468 | XM_021616793.1 | 9 | 12a | Yes |
| acvr1ba2 | LOC110491966 | XP_021421545 | XM_021565870.1 | 16 | 12a | Yes |
| acvr1bb1 | LOC110494313 | XP_021424945 | XM_021569270.1 | 17 | 12b | Yes |
| acvr1bb2 | LOC110528117 | XP_021465613 | XM_021609938.1 | 7 | 12b | Yes |
| acvr1ca1 | LOC110501520 | XP_021434837 | XM_021579162 | 22 | 3a | Yes |
| acvr1ca2 | LOC110520051 | XP_036830206 | XM_036974311.1 | 3 | 3a | Yes |
| acvr2aa1 | LOC110520186 | XP_021453003 | XM_021597328.1 | 3 | 3a | Yes |
| acvr2aa2 | LOC110501883 | XP_021435411 | XM_021579736.2 | 22 | 3a | Yes |
| acvr2ab1 | LOC110495930 | XP_021427133 | XM_021571458.2 | 18 | 13b | Yes |
| acvr2ab2 | LOC110527542 | XP_036838921 | XM_036983026.1 | 7 | 13b | Yes |
| acvr2ba1 | LOC100301655 | NP_001177870 | NM_001190941.1 | 15 | 13a* | Yes<sup>#1</sup> |
| acvr2ba2 | LOC110535274 | XP_021475849 | XM_021620174.1 | 11 | 13a | Yes |
| acvr2bb | LOC110531045 | XP_036842531 | XM_036986636.1 | 8 | 13b | Yes |
| bambia1 | LOC100301651 | NP_001153956 | NM_001160484.1 | 11 | 13a* | Yes |
| bambia2 | LOC110490039 | XP_036800238 | XM_036944343.1 | 15 | 13a | Yes |
| fstb1 | LOC110537003 | XP_021478374 | XM_021622699.2 | 12 | 9b | No |
| fstb2 | LOC110523922 | XP_036833533 | XM_036977638.1 | 5 | 9b | No |
| fsta1 | LOC100301657 | NP_001153960 | NM_001160488.1 | 11 | 9a* | Yes<sup>#2</sup> |
| fsta2 | LOC100301646 | NP_001153955 | NM_001160483.1 | 6 | 9a | Yes<sup>#2</sup> |
| gdf11b1 | LOC110494206 | XP_021424716 | XM_021569041.2 | 17 | 12b | Yes |
| gdf11b2 | LOC110528233 | XP_021465784 | XM_021610109.2 | 7 | 12b | Yes |
| inhab1 | LOC100135804 | NP_001117672 | NM_001124200.1 | 18 | 3b* | Yes<sup>#3</sup> |
| inhab2 | LOC110528842 | XP_021466674 | XM_021610999.1 | 7 | 3b | Yes |
| inhbaa1 | LOC110490008 | XP_021418759 | XM_021563084.2 | 15 | 13a | Yes |
| inhbaa2 | LOC110535572 | XP_021476318 | XM_021620643.1 | 11 | 13a | Yes |
| inhbab1 | LOC110509121 | XP_021445767 | XM_021590092.1 | 28 | 13b | Yes |
| inhbab2 | LOC110529989 | XP_021468383 | XM_021612708.1 | 8 | 13b | Yes |
| inhbba1 | LOC110520286 | XP_021453151 | XM_021597476.1 | 3 | 3a | Yes |
| inhbba2 | LOC110501797 | XP_021435309 | XM_021579634.1 | 22 | 3a | Yes |
| inhbbb1 | LOC110527500 | XP_021464487 | XM_021608812.1 | 7 | 3b | Yes |
| inhbbb2 | LOC110495886 | XP_021427073 | XM_021571398.1 | 18 | 3b | Yes |
| inhbcb1 | LOC110494776 | XP_021425781 | XM_021570106 | 17 | 12b | Yes |
| inhbcb2 | LOC110527696 | XP_021464812 | XM_021609137.1 | 7 | 12b | Yes |
| mstna1 | LOC100135919 | NP_001117754 | NM_001124282.1 | 22 | 3a* | Yes<sup>#4</sup> |
| mstna2 | LOC100135920 | NP_001117755 | NM_001124283.2 | 3 | 3a* | Yes<sup>#4</sup> |
| mstnb1 | LOC100136340 | NP_001118048 | NM_001124576.1 | 7 | 3b* | Yes |
| mstnb2 | LOC110496923 |  | XR_002469620 | 18 | 3b* | Yes |
| smad2a1 | LOC101867532 | NP_001268237 | NM_001281308.1 | 11 | 9a | Yes<sup>#5</sup> |
| smad2a2 | LOC101268919 | NP_001268263 | NM_001281334.1 | 6 | 9a | Yes<sup>#5</sup> |
| smad3a1 | LOC110526791 | XP_036837330 | XM_036981435.1 | 6 | * | Yes |
| smad3a2 | LOC100305196 | NP_001158572 | NM_001165100.1 | 26 | * | Yes |
| smad3b | LOC118945776 | XP_036824526 | XM_036968631.1 | 30 | * | No |
| smad4ab1 | LOC110537152 | XP_036793045 | XM_036937150.1 | 12 | 9b* | No |
| smad4ab2 | LOC110523770 | XP_021458448 | XM_021602773.2 | 5 | 9b | Yes |
| smad4aa1 | LOC110525240 | XP_036835280 | XM_036979385.1 | 6 | 9a | Yes |
| smad4aa2 | LOC101268920 | XP_021476516 | XM_021620841.2 | 11 | 9a | Yes |
| smad4bb1 | LOC101268921 | XP_021413475 | XM_021557800.2 | 2 | 2b | Yes |
| smad4bb2 | LOC110503878 | XP_036827746 | XM_036971851.1 | 3 | 2b | Yes |
| smad4ba1 | LOC110488280 | XP_036826852 | XM_036970957.1 | 14 | 2a | Yes |
| smad4ba2 | LOC110496821 | XP_036808764 | XM_036952869.1 | 18 | 2a | Yes |
| smad6a1 | LOC110509922 | XP_021446816 | XM_021591141.2 | 29 | 10a* | Yes |
| smad6a2 | LOC118936600 | XP_036819676 | XM_036963781.1 | 26 | 10a* | No |
| smad6b | LOC110510499 | XP_036824722 | XM_036968827.1 | 30 | 10b* | No |
| smad7b1 | LOC101268922 | NP_001268265 | NM_001281336.1 | 5 | 9b* | No |
| smad7b2 | LOC110536948 | XP_036792774 | XM_036936879.1 | 12 | 9b | Yes |

\* denotes genes not included in the Parey et al. 2022 ancestral chromosome tracking. Putative ancestral chromosomes identified based on conserved chromosomes from other ohnologs.

Northern Pike (*E. lucius*) are a close relative to the salmonid lineage and do not possess the Ss4R WGD but do possess the Ts3R genome duplication (Lien et al. 2016; Davesne et al. 2021).

Peptide sequences were used to calculate the evolutionary distance between ohnologs as a measure of the level of divergence between Ts3R and Ss4R duplication events (Figure 1A). As expected, genes separated by the Ts3R duplication event exhibited higher divergence in peptide sequence reflecting the longer evolutionary time from which that duplication event occurred (Figure 1A). However, some duplicated members of the pathway remained remarkably conserved across both genome duplication events (e.g., Smad signal transduction molecules *smad2, smad3*, and *smad4;* Figure 1A), suggesting stronger selection pressures on specific pathway genes to maintain sequence integrity. The Ss4R ohnologs from singleton Ts3R genes (i.e., single copy genes in Northern Pike) were not more likely to exhibit peptide sequence divergence as Ss4R ohnologs than four copy gene families (p=0.95). This phenomenon has previously been identified across the RBT genome showing that low sequence divergence of Ss4R ohnologs is not associated with gene retention (Berthelot et al. 2014).

**Figure 1.**
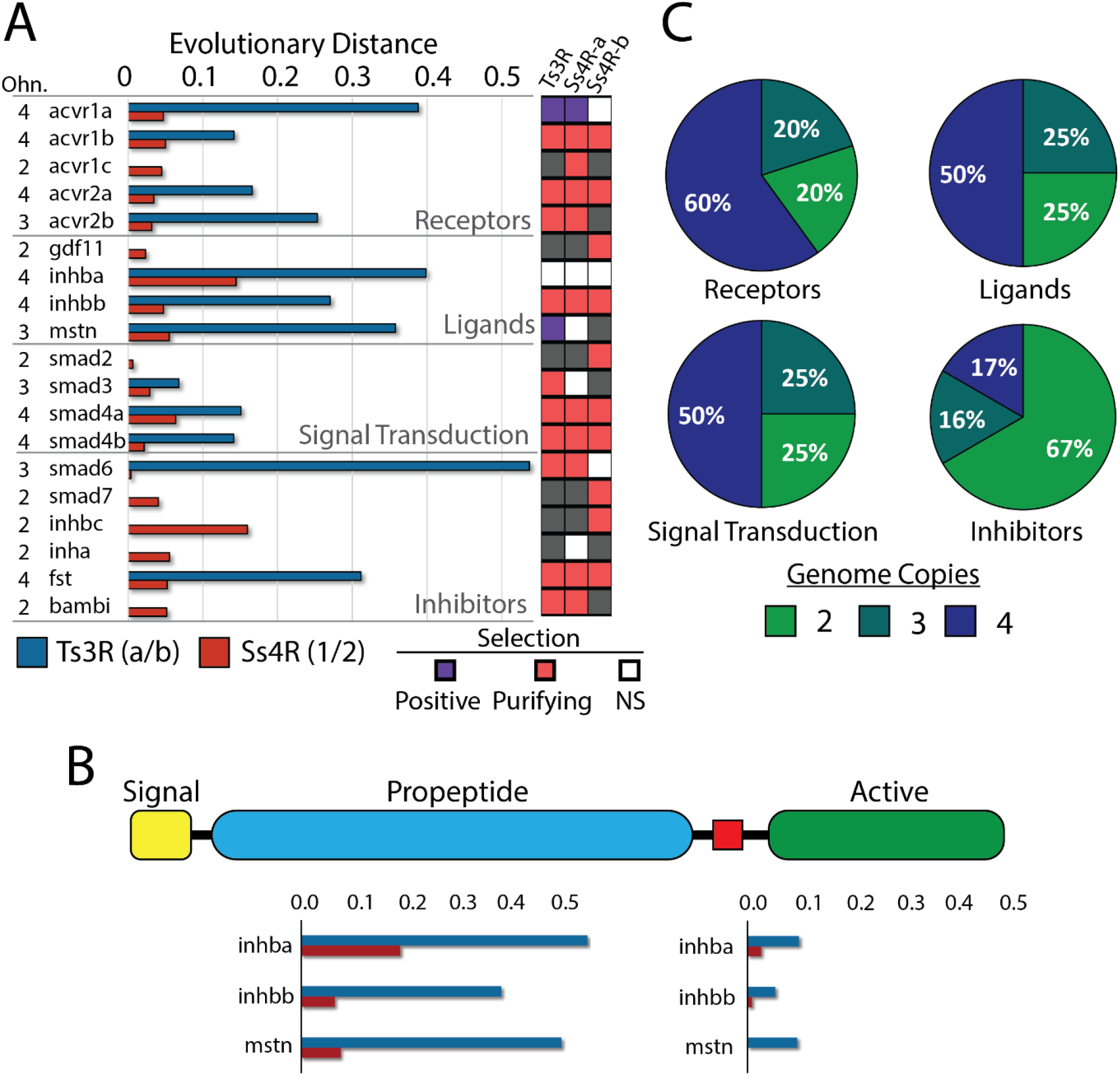
Evolutionary distance and ohnolog retention following whole genome duplication of the activin receptor signaling pathway. **(A)** Evolutionary distances between Ts3R (blue) and Ss4R (red) ohnologs along with their signatures of selection. Positive (purple), purifying (red), and no significance (NS; white) in selection is identified (gray; no ohnolog comparison possible). **(B)** Characteristic protein domains of select activin pathway ligands and the difference in evolutionary distance between the propeptide and active domains of the proteins. **(C)** Percentage of ohnolog copy numbers from the four main activin pathway signaling components.

Given the close relationship between the activin receptor signaling pathway genes in RBT, it was not surprising that signatures of purifying selection were observed for the majority of the ohnologs (Figure 1A). Evidence of potential positive selection pressure was observed for the activin receptor 1a (*acvr1a*) and myostatin gene families (*mstn*; Figure 1A). Myostatin is a major muscle growth regulator in mammals and fish but is found to be expressed in many more tissues in fish, suggesting possible alternative functions (McPherron and Lee 1997; Garikipati et al. 2006; Kim et al. 2019).

Smad signal transduction ohnologs were especially conserved across both WGD events. The peptide sequence of Smad2a1 and Smad2a2 for example exhibit 99% sequence similarity.

Smad3 is unique in that the family consists of three genes, two ohnologs that are related to *E. Lucius smad3a*, and a single gene related to *E. Lucius smad3b*. Interestingly, *O. mykiss smad3a2* is severely truncated such that it only produces a fully intact Smad MH1 domain but lacks the characteristic Smad MH2 domain (Figure S1). The MH1 domains of Smad3a1 and Smad3a2 are highly conserved, exhibiting a 97% sequence similarity (2 aa difference out of 70). Smad MH2 domains interact with and are phosphorylated by type I serine-threonine kinase receptors to activate the signaling pathway. They are also required for homodimer formation with other Smad3 proteins or heterodimer formation with Smad4, the common partner Smad (Co-Smad) essential for signal transduction (Moustakas 2002). The MH1 domain of Smad proteins is important for binding to Smad-Binding Elements (SBE) on the DNA during the activation of downstream gene transcription. When bound to DNA the MH2 domain of Smad proteins is involved in the recruitment of transcription factors, co-activators, and co-repressors. While the functional significance of the truncated *smad3a2* ohnolog containing only a MH1 domain in RBT is unknown, previous research has shown that experimental truncation of Smad3, to remove the c-terminal MH2 domain, produces a dominant negative signaling molecule that can interfere with Smad signal transduction (Liu et al. 2001). Given the high sequence similarity between the *Smad3a* ohnologs in RBT, it is unlikely that *smad3a2* is a non-functional pseudogene since such high purifying selection would not be expected for a pseudogene lacking a cellular function (*smad3a1* and *smad3a2* have 96.8% nucleotide sequence identity).

The majority of activin receptor signaling pathway ligands exhibited high peptide sequence distance between Ts3R ohnologs, some of the highest of all activin signaling pathway ohnologs (e.g., *inhba, inhbb, mstn*; Figure 1A). However, these differences were primarily found within the propeptide domain of the molecule, with high conservation of the active domain of the peptides (Figure 1B). TGF-β ligands are post-translationally processed to remove the propeptide domain from the active domain of the peptide. The propeptide domain can then function as an inhibitor of the active dimer adding an additional regulatory mechanism to the signaling cascade (Constam 2014). As was observed in RBT, there is often high sequence conservation between the active domain of TGF-β ligands, even across distant taxa (Rodgers et al. 2007). The variation in the propeptide domains suggests the presence of differential selective pressures that exist on the inhibitory propeptide domain vs. the active domain of activin receptor signaling pathway ligands. Alternatively, it is also possible that propeptide and active domain binding may only require a few conserved regions to function, allowing sequence divergence to occur throughout the propeptide domain without altering its activity. The propeptide of activin A is known to have low affinity to the mature activin A molecule and there is sequence conservation in key regions of TGF-β, BMP, and activin A propeptide domains despite otherwise being unrelated (Wang et al. 2016).

The TGF-β superfamily commonly retains duplicated members following WGD events, giving rise to the extended family of genes found in most vertebrate species (Zheng et al. 2018). All of the 19-core activin receptor signaling pathway gene families that we analyzed in RBT contained at least 2 duplicated copies with 63% of the genes exhibiting three or more copies (Figure 1C). Four copy gene families were common for receptors, ligands, and Smad4 but uncommon for inhibitory proteins and R-Smads (Figure 1C). Smad signal transduction molecules were distinct in that both of the R-Smads, Smad2 and Smad3, have effectively reverted to two gene copies in RBT (note the truncation of *smad3a2* described earlier). The Co-Smad, *Smad4*, however, exhibits the highest number of activin receptor signaling pathway ohnologs, with the family consisting of eight highly conserved members (Table 1, Figure 1A). This suggests that the loss of *smad4* family members is not tolerated despite the opposite selective pressure at work for R-Smad proteins. *Smad4* is unique among Smad signal transduction molecules as it is the sole Co-Smad for the Smad signaling cascade, and complexes with R-Smad proteins to facilitate gene transcription. While R-Smads differ between the various TGF-β pathways, all TGF-β pathways utilize Smad4. Its central role in multiple TGF-β superfamily pathways may function as a bottleneck on evolutionary diversification since alterations in the gene family members could influence a wide range of physiological processes. All *smad4* family members in RBT exhibit high peptide sequence conservation in the MH1 and MH2 domains with increased sequence divergence in the linker domain (Figure S2). This is similar to what has been observed in Japanese flounder (Yu et al. 2020). In this model system, evidence suggests that structural changes in the linker region may influence Smad4 protein function despite having highly conserved MH1 and MH2 domains (Yu et al. 2020). If gene dosage is the primary driver of smad4 ohnolog retention we could expect to observe partitioning of the expression between ohnologs such that members would either share expression (hypofunctionalization) or exhibit expression specialization to specific tissues (subfunctionalization). An alternative scenario could be related to the production of toxic *smad4* gene products since Smad4 is a known tumor suppressor who’s loss of function can increase susceptibility to several forms of cancer (Takaku et al. 1999; Wang et al. 2007).

An extreme example of Ts3R ohnolog divergence was noted for the *smad6* gene family, which consists of three extant family members (Figure 1A). The s*mad6a* gene family in RBT has a highly conserved sequence similarity, one of the highest across all of the gene families analyzed, whereas the *smad6b* ohnolog pair has lost a member and has evolved a substantially different peptide sequence compared to *smad6a* (Figure 1A). Smad6 functions along with Smad7 as an inhibitor of the Smad signaling cascade. The *smad6* gene family is one of only two activin receptor signaling pathway inhibitors that remained duplicated after the Ts3R WGD and it is possible that the observed sequence divergence may reflect a change in gene function of *smad6b* in RBT.

### Expression Distribution of Activin Receptor Signaling Pathway Ohnologs

In addition to coding sequence divergence, functional evolution of ohnologs can occur at the level of gene regulation, by altering the tissue distribution or level of gene expression. To investigate potential regulatory functionalization of the activin receptor signaling pathway ohnologs, we employed NanoString nCounter technology to provide a quantitative measure of gene expression for 49 members of the activin receptor signaling pathway across 23 adult RBT tissues (Figure 2A, and S3). This included 7 distinct skeletal muscle regions or specialized muscle groups (e.g., extraocular muscle, adductor muscle, etc.) due to the importance of the pathway in skeletal muscle growth and development (Figure S3). The genes selected for gene expression profiling represented members of all 19 activin signaling pathway gene families along with most duplicated ohnologs (Table 1 and S1).

**Figure 2.**
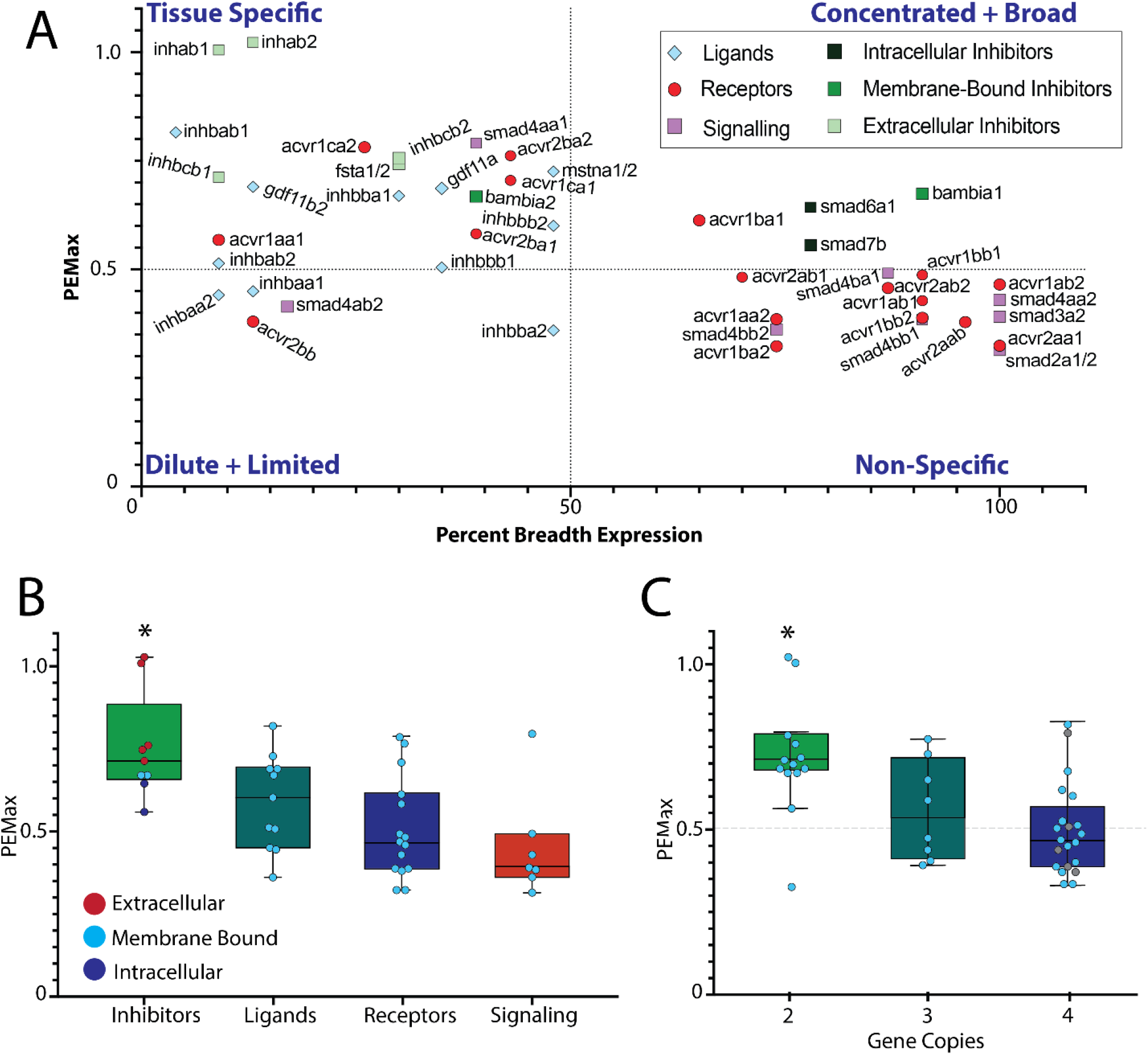
Tissue specificity of activin receptor signaling pathway ohnologs. **(A)** The tissue specificity of each gene was determined by both the percent breadth of expression and the PEM_MAX_. Based on these values, the overall pattern of expression was categorized into quadrants of tissue specificity based on the strength of each measure. **(B)** Quantification of tissue specificity between activin pathway components. **(C)** Analysis of the relationship between tissue specificity and ploidy status for activin signaling pathway ohnologs. *Denotes significant differences between inhibitors (B) and 2 copy ohnolog families (A) compared to other groups (p < 0.05).

Tissue specificity was calculated for each gene using two measures of specificity: percent breadth of expression and Preferential Expression Measure (PEMMAX). The percent breadth of expression is a measure of tissue specificity independent of the expression in other tissues, while the PEM_MAX_ measures the extent to which the gene’s transcription profile is concentrated into one tissue (Huminecki et al. 2003). PEM_MAX_ helps account for genes that may be broadly expressed across a wide range of tissues at low levels, but which have high levels of expression in a small amount of tissues. The combination of the two calculations was used to provide a detailed analysis of the type of expression pattern observed by each gene in the pathway by categorizing the pattern into tissue-specific, non-tissue specific, concentrated but broadly expressed, or dilute with limited tissue expression, based on the strengths of each tissue specificity measure (i.e., breadth of expression and PEM_MAX_; Figure 2A). The gene families were analyzed according to their assumed function to identify trends in specific signaling factor components.

When analyzing across functional groups, inhibitors exhibited the most tissue-specific expression profile (Figure 2A and 2B). Sixty-seven percent of inhibitors (6 out of 9) were expressed in a tissue-specific manner (Figure 2A). The inhibitors that were expressed across a wide range of tissues were still expressed at elevated levels in specific tissues (i.e., concentrated but broadly expressed; Figure 2A). The ligands of the activin receptor signaling pathway also showed a tissue-specific expression profile with 73% exhibiting a tissue-specific expression pattern with the remaining having expression limited to a low number of tissues (Figure 2A). There was no identifiable pattern distinguishing between the tissue specificity of Type I vs. Type II receptors but overall, 33% of receptors exhibited a tissue-specific expression profile and 67% were non-specific (Figure 2A). An overall broad expression pattern was also observed for Smad signal transduction molecules which only had one gene expressed selectively in skeletal muscle tissues (*smad4aa1*), and 71% of genes had non-specific expression (Figure 2A). The expression pattern of *smad4aa1* is of note since it does not fit the expression pattern of other Smad4 proteins, which are expressed across a wide range of tissues, reflective of their role as Co-Smads for all TGF-β pathways. The divergent expression of *smad4aa1* is potential evidence of regulatory specialization of this Co-Smad in skeletal muscle.

Quantification of the level of tissue specificity of activin receptor signaling pathway functional groups was performed using the PEM_MAX_ values for each gene as previous work suggests that this measure may be more representative of tissue specificity for ohnologs than analyzing the breadth of tissue expression (Huminiecki et al. 2003). On average, inhibitors were identified as the most tissue-specific group of molecules (0.75 ± 0.053), with signaling transducers the least tissue-specific (0.45 ± 0.060; p=0.002; Figure 2B). Inhibitors can interact with the signaling pathway as extracellular, membrane-bound, and intracellular regulators. To determine if the cellular location of inhibitors influenced their specificity, we independently analyzed inhibitors based on their location of influence (Figure 2B). While extracellular inhibitors trended toward being more tissues specific (Figure 2B), there were no significant differences in tissue specificity between the classes of inhibitors. It is important to note, however, that *smad6* and *smad7* were expressed at low levels across a large number of tissues which is unlike most of the other pathway inhibitors. This is not surprising as these genes have roles in regulating the signal transduction of other TGF-β signaling pathways which likely requires a broader expression pattern than activin receptor signaling pathway-specific inhibitors.

The relationship between the expression pattern of activin pathway functional groups and the ploidy status of the ohnologs was analyzed to look for signatures of specialization to different tissues and whether this might have contributed to the retention of ohnologs after WGD events. We directly compared the tissue specificity of 2, 3 and 4 copy gene families and discovered that the 2 copy gene families were more likely to be expressed in specific tissues compared to both 3 (p=0.049), and 4 (p=0.0004) copy gene families (Figure 2C). Even without the influence of the *smad4* gene family, which contains numerous family members that are expressed broadly across tissues, 4 copy gene families are significantly more non-specific than 2 copy families. Overall, there is a clear association between the retention of ohnologs following WGD and their expression pattern across tissues (Figure 2C). However, this relationship is not universal. While 2 copy gene families, like most inhibitors, are likely to exhibit tissue-specific expression, having tissue-specific expression does not necessarily drive the loss of ohnolog members, since most ligand families are tissue-specific but contain many fully duplicated gene families (Figure 1A). The relationship between ohnolog retention and expression profile appears to be more heavily influenced by gene function than whether genes exhibit a tissue-specific expression pattern.

### Functionalization themes across the Ts3R and Ss4R whole genome duplication events

A leading theory to explain the fate of duplicated genes following WGD is that pressure to maintain optimal gene dosages for dosage-dependent pathways influences the retention of ohnologs (Gillard et al. 2021; Birchler and Yang 2022). To maintain balance, these genes can alter their expression dynamics such that they either reduce the expression of each ohnolog to meet the overall dosage requirement of the cell or they can partition their expression between cell types that were originally expressed by the ancestral duplicated gene. Using the twin duplication events in salmonids as a model, we investigated the mechanism by which ohnologs modulate their expression post-WGD. A classification system was used to categorize the levels of gene expression into 1) Not expressed, 2) Low Expression, 3) Moderate expression, and 4) High expression genes (Figure 3A). This analysis was performed on gene from which expression data was available for all of the ohnologs in the gene family. Categorizing gene expression allowed for standardization of gene expression levels to account for the influence of extremely highly expressed genes as well as to enable unambiguous comparison between the expression classifications.

**Figure 3.**
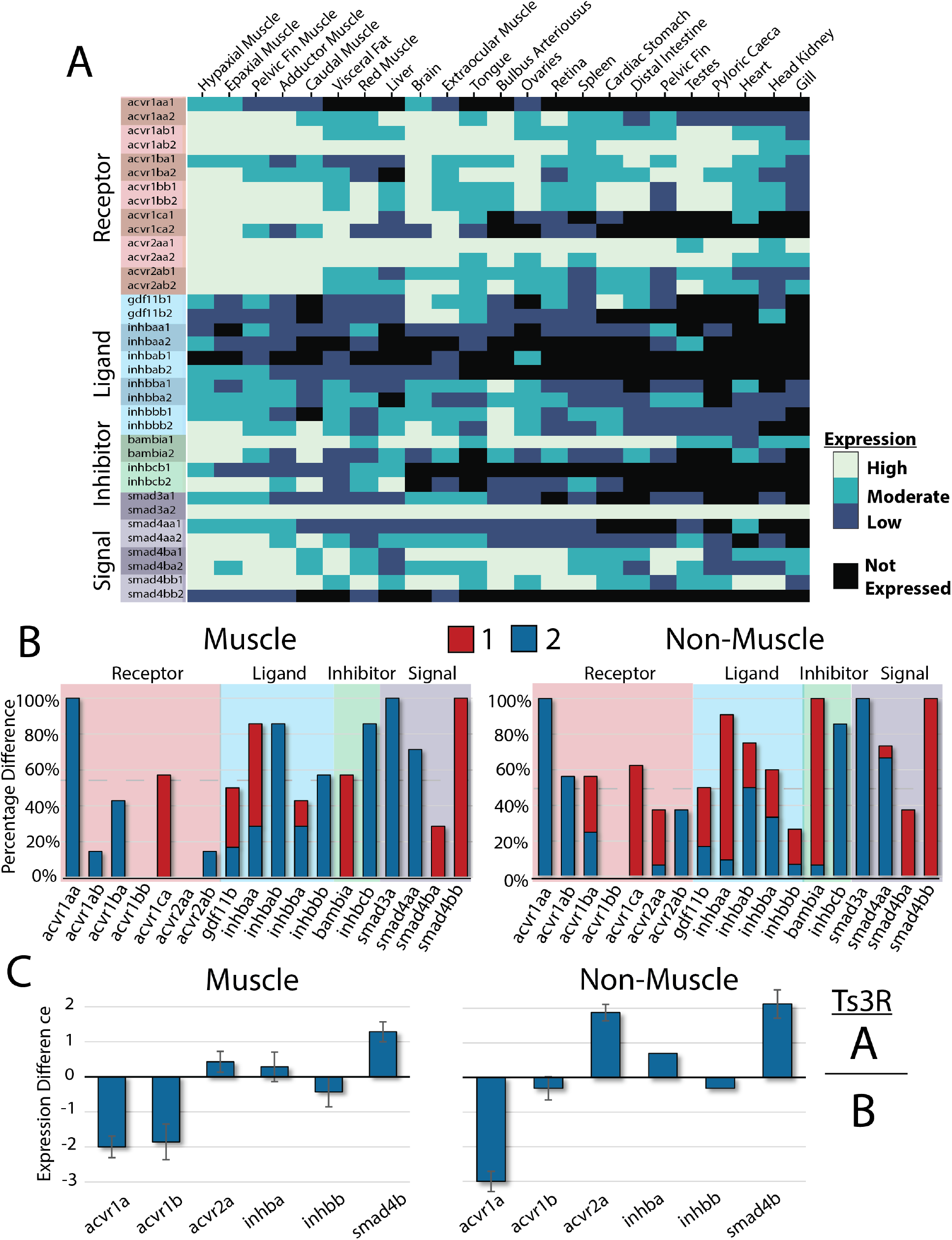
Regulation of activin pathway ohnolog expression patterns across Ts3R and Ss4R whole genome duplication events. **(A)** Expression heat map categorizing low, medium, and high levels of expression for each gene across tissues. Molecule types are identified for receptors (red), ligands (blue), inhibitors (green), and signal transduction molecules (purple). **(B)** The difference in expression of specific Ss4R ohnologs (1 vs. 2) as a percentage of the tissues the family is expressed in. A low percentage represents common levels of expression across ohnologs while high percentages reflect a preference for a dominant form expressed in most tissues. The selective expression of different dominant forms (1-red vs. 2-blue) in select tissues is identified by multi-color bars. **(C)** The average difference in expression between Ts3R ohnolog pairs (a vs b). Positive values represent preference in the expression of the Ts3R-a family while negative values show a dominance of the Ts3R-b family.

The level of gene expression between ohnologs was examined by calculating the difference in gene expression between each Ss4R ohnolog pair in tissues in which one or both of the genes were expressed. The pattern of dominance was heavily dependent on the molecule type. For inhibitors and Smad signal transduction molecules, a single Ss4R ohnolog was usually observed to be dominantly expressed across most of the tissues (Figure 3B). This was surprising for Smad signal transducton molecules since they are expressed across a wide range of tissues and exhibit both 2 copy and 4 copy gene families. They also exhibit high levels of coding sequence conservation and evidence of purifying selection against protein coding changes (Figure 1A), yet we identified divergence in the level of gene expression between ohnologs (Figure 1A). The inhibitors that were included in the analysis were all from 2 copy gene families known to exhibit a tissue specific expression pattern. These findings suggest that partitioning of gene expression across tissues (subfunctionalization) is not a driving mechanism for inhibitors and signal transduction molecule but instead regulatory suppression of one ohnolog over the other predominates. The preference for a single Ss4R ohnolog to be prioritized in tissues despite both ohnologs being expressed in most tissues could represent a form of hypofunctionalization where dosage is maintained primarily by a single gene.

For some ohnolog pairs, it is possible that suppression of the expression of a single form may help enhance the pseudogenation process and that the expression pattern that we observe is a hallmark of the eventual loss of the low expressing gene copy. An example of this can be observed with the inhibitor activin Cb (*inhbcb*). Of the activin Cb ohnologs, both *inhbcb2* and *inhbcb1* have the same expression profile (expression in skeletal muscle and liver), however, *inhbcb2* is expressed significantly higher than *inhbcb1* in all tissues (p = 0.014; Figure S4A). The evolutionary distance between *inhbcb1* and *inhbcb2* is also the second largest amongst Ss4R ohnologs (0.1598). In mammals, activin C is believed to be an inhibitor of activin A and is expressed primarily in the liver, however, its function in that tissue is not fully understood (Gold et al. 2009). In addition to liver tissue, we identified high expression of activin C in skeletal muscle tissue in RBT, suggesting a potential secondary function for activin C in fish. The expression pattern of activin C ohnologs provides evidence of regulatory suppression of *inhbcb1* in RBT, which may help drive the future pseudogenization of this ohnolog, as previously occurred after the Ts3R WGD.

In contrast to inhibitors and Smad signal transduction molecules, activin signaling pathway ligands commonly had independently dominant expression patterns between the ohnologs (Figure 3B). This partitioning of expression may represent a mechanism by which the cell is able to differentially regulate the activity of the pathway across tissues. Activin Ab (*inhbab1* and *inhbab2*) is a putative example of this subfunctionalization of the activin receptor signaling pathway ligands. Activin Ab ohnologs have the highest evolutionary distance amongst all activin receptor signaling pathway Ss4R ohnologs (0.1929; Figure 1B). Unlike activin C however, there is not a single dominantly expressed ohnolog, instead, *inhbab* genes exhibit differential expression in skeletal muscle (*inhbab2*) and ovarian (*inhbab1*) tissue (Figure S4B). Activin A is known to be commonly expressed in these tissues in vertebrates since activin A is a multifunctional signaling molecule that has key roles in reproductive tissues (Hsueh et al. 1987), skeletal muscle (Latres et al. 2017), and embryonic development (Vassalli et al. 1994). The expression profile of activin Ab suggests that in RBT, the activin Ab ohnologs have subfunctionalized to independently accomplish the cellular functions that are carried out by a single gene in other vertebrate species.

Despite observing a variety of expression patterns, receptor ohnologs were more likely to share expression equally across tissues than the other signaling pathway components (Figure 3A and 3B). Activin receptor 1Bb ohnologs (*acvr1bb*) were an extreme example of this shared expression, with both ohnologs being expressed at high, equal levels across all of the analyzed tissues (Figure 3A). Unlike activin receptor 1Bb, activin receptor 1Aa (*acvr1aa*) had a single ohnolog predominate across all tissues, and dominant expression was equally partitioned between the ohnologs of activin receptor 1Ba (*acvr1ba*; Figure 3B). It is important to note that activin receptor 1Aa was also the only activin receptor ohnolog pair that exhibited a signature of positive selection (Figure 1A). It is clear from our tissue distribution (Figure 2) and expression level characterization that activin receptors are relatively ubiquitous across tissues and that changes in the expression of receptors is likely not an important regulator of activin signaling pathway. It is possible that the pressure to ensure a sustained expression of receptors (gene dosage) is a driving mechanism for the high rate of ohnolog retention in these molecules, but this remains to be experimentally validated.

In order to gain insight into the long-term evolutionary functionalization of duplicated ohnologs an understanding of the ancestral state of expression is needed. This is challenging given that gene regulatory mechanisms can change in unknown ways over evolutionary time, which can influence the interpretation of ancestral gene function. Under the putative framework that gene dosage requirements will drive the overall expression level of ohnologs, the gene expression dynamics of the ancestral Ts3R ohnolog might be estimated through the combined expression of duplicated Ss4R ohnologs. We employed this approach to gain insight into how members of the activin receptor signaling pathway may have altered their expression pattern in response to genome duplication. The RBT model system and its two WGD events facilitated this analysis within a single species, which has advantages over making cross-species comparisons that are influenced by species-specific differences (e.g., body plan, life history, etc.), which can reduce confidence in accurately establishing the ancestral gene expression patterns.

The estimated ancestral Ts3R expression dynamics were analyzed using the total expression level of Ss4R ohnologs in fully tetraploid gene families (Figure 3C). The activin receptor 1Ab was dominantly expressed over 1Aa ohnologs, which is similar to the pattern of dominance identified for specific Ss4R ohnologs of that gene family (1 vs. 2; Figure 3B). *Smad4Bb* also showed a pattern of dominance over the s*mad4Ba* ohnolog pair that reflects the dominance observed for *Smad4Bb1* Ss4R ohnologs (Figure 3C). The Ts3R ligand ohnologs also exhibited a similar overall pattern to that of Ss4R ligand ohnologs, suggesting that subfunctionalization is a common mechanism of ohnolog retention for activin receptor signaling factor ligands (Figure 3C). In contrast to the expression of Ss4R ohnologs, the Ts3R activin receptor 2Aa ohnolog pair (*acvr2a*), a type II receptor, was dominantly expressed in most non-muscle tissues something that was not seen in either of the Ss4R ohnolog families (i.e., *acvr2aa1/2* and *acvr2ab1/2*). Besides this outlier, the majority of Ts3R gene families exhibited similar expression patterns to the more recent Ss4R gene families. This provides evidences suggesting that the core function of specific genes within a signaling pathway has an important role in determining the fate of ohnologs following WGD events.

While we only analyzed genes involved in the activin receptor signaling pathway in this study, it is likely that gene function has an important role in other signaling pathways or interconnected gene networks. Further research on the influence of gene function on ohnolog evolution will provide needed insights into the molecular pressures influencing ohnolog evolution following whole gene duplication, and its role in shaping the genome of organisms.

## Materials and Methods

### Animal Care and Use

The rainbow trout used in this study were housed in a centralized facility at Washington State University. All experiments were performed in accordance with preapproved protocols reviewed by the Institutional Animal Care and Use Committee of Washington State University (Protocol #6607). To reduce the influence of genetic variation in our activin pathway gene expression atlas we utilized YY male clonal RBT from the Swanson line, for which the first draft of the RBT genome sequence was derived (Berthelot et al. 2014). Ovarian tissue samples were obtained from female clonal XX RBT of the Oregon State line (Young et al. 1998). Clonal RBT are genetically identical homozygous fish created through androgenesis (YY) or gynogenesis (XX). The use of clonal RBT in establishing the activin receptor signaling pathway gene expression atlas helped to minimize expression variation that may be impacted by genetic variation between individuals.

### Tissue Collection

Twenty-three tissues were extracted in triplicate from 2-3 year old clonal adult RBT (Figure S3). All tissue samples were collected from a minimum of three individuals, flash-frozen in liquid nitrogen and stored at -80ºC until total RNA extraction.

### RNA Preparation and NanoString Analysis

RNA was extracted from the tissue samples using either TRIzol or RNA extraction kits from Zymo and Qiagen. Optimal extraction methods to obtain high RNA yields and quality differed for each tissue. Purified RNA was quantified on a Qubit fluorometer and quality was assessed using a 5200 fragment analyzer performed by the WSU Laboratory for Biotechnology and Bioanalysis (LBB) core. The resulting RNA was sent to the Fred Hutchinson Cancer Research Center, Genomics and Bioinformatics core for Nanostring nCounter analysis. The custom NanoSting nCounter probe set consisted of 49 of the 59 analyzed RBT activin signaling pathway genes/ohnologs and three reference genes (ef1a, tbp, and actb; Probe Sequences found in Table S1). All data received from Nanostring was normalized in the nSolver program using *ef1a, tbp*, and *actb* as reference genes under default settings.

### Molecular Phylogeny

Protein coding sequences for the analyzed activin pathway genes were obtained for both RBT (*Oncorhynchus mykiss;* USDA_OmykA_1.1) and a closely related non-salmonid species, Northern Pike (*Esox lucius;* fEsoLuc1.pri). To identify likely ohnologs, ORTHOSCOPE software was used to compare the protein-coding sequences of each RBT gene family to that of *E. lucius*, creating the gene family trees used for naming (Inoue and Satoh 2018). Suggested gene names were given based on standard gene naming conventions with the Ts3R duplicated ohnologs identified alphabetically based on the derived ancestral chromosomes and Ss4R duplicated ohnologs designated numerically (Parey et al. 2022). Previous nomenclature of the RBT myostatin gene followed different gene naming conventions with the Ss4R duplicated ohnologs identified alphabetically and Ts3R duplicated ohnologs designated numerically (Rodgers et al. 2007). The new proposed gene names reflect the inclusion of ancestral teleost chromosomes of origins as well as current standard vertebrate nomenclature conventions. All of the activin pathway gene information and proposed nomenclature can be found on Table 1. The pairwise distance and selection analysis was calculated for each gene family using MEGA X software (Kumar et al. 2018) using a Poisson model and Nei-Gojobori method, respectively.

### Determining Tissue Specificity

To characterize the tissue specificity patterns within and across ohnolog gene families, a conservative low abundance threshold of expression was employed to identify the primary genes expressed in specific tissues above the identified baseline level of expression. This low abundance threshold for expression was determined by calculating the 50th quartile of the normalized expression counts for each molecule type across all tissues individually (receptors = 87, inhibitors = 36, ligands = 26, signal transducers = 100), since each molecule type exhibited variability in their baseline expression level (i.e., inhibitors and ligands, on average, are much lower expressed than receptors and signal transducers). Percent breadth of expression was calculated as the percentage of tissues (16 non-muscle and 2 skeletal muscle) expressing the ohnolog above low abundance levels. The Preferential Expression Measure (PEM) was defined as log10(S/A) such that S is the specific expression for a single gene in one tissue and A is the mean gene expression across all tissues. The PEM_MAX_ is the maximal value of the PEM (Huminiecki et al. 2003). To improve the interpretation of tissue specificity for individual ohnologs, the PEM_MAX_ was directly compared to the percent breadth of expression for each gene. In order to not bias the results to muscle tissue, since multiple muscle regions were sampled (n=7), the analysis of tissue specificity was only analyzed using two regions of epaxial skeletal muscle. Tissue specificity of the genes were grouped by functional group (ligand, receptor, inhibitor, signaling protein) and the PEM_MAX_ was employed to determine significant differences between molecule groups and duplication status (Figure 2B and 2C).

### Ohnolog Expression Differences

Characterization of differential regulation between ohnologs was performed by categorizing expression levels into defined levels of expression to adjust for extremes (ranging from 0.00 to 2104.77) that can interfere with making direct comparisons between ohnolgs. The overall expression data for all genes was classified into quartiles (0 to 18.25, 18.26 to 44.42, 44.43 to 138.76, and 138.77 to 2104.77) and each gene was assigned a value of 0, 1, 2, or 3 based on the level of its expression. This categorized gene expression into the following four groups: not expressed, low expression, medium expression, and high expression (Figure 3). The expression categories were assigned for each tissue and the difference in ohnolog expression was calculated for both Ts3R (a/b) and Ss4R (1/2) ohnologs (Figure 3B and 3A).

### Statistical Analysis

A one-way ANOVA was performed to determine differences across molecule types followed by post hoc Tukey tests to identify significant differences within the ANOVA results. Paired t-tests were used to determine pairwise differences in ohnolog expression between specific groups when necessary. The p value was set at <0.05.

## Data and Resource Availability

The raw NanoString nCounter gene expression data used in the study can be found in the NCBI Gene Expression Omnibus data archive and at https://mphelpslab.org/resources.

## Acknowledgements

The concepts and experimental designs of the research were produced by J.R. and M.P. The experimental procedures were performed by J.R. with help on data analysis by M.P. and L.D. The manuscript was written by J.R. and M.P. The research was supported by a USDA National Institute of Food and Agriculture grant # 2021-67015-33400.

## Figure Legends

**Figure S1. Smad3 truncation and alignment.** Diagram of the MH domains of RBT Smad3 peptide ohnologs showing truncation of Smad3a2 to contain only the MH1 domain. A multiple sequence alignment shows high conservation of the smad3 ohnologs.

**Figure S2. Variability in linker regions of Smad4 peptide sequences.** The peptide sequence alignment of Smad4a demonstrates the high sequence conservation between ohnologs at the MH1 (yellow) and MH2 (orange) domains with variation found in the linker region of the peptide sequence.

**Figure S3. Tissue sampling regions.** The diagram represents the 23 tissues used for gene expression analysis of the activin receptor signaling pathway, including 7 distinct muscle regions.

**Figure S4. Examples of hypofunctionalization and subfunctionalization of *inhbcb* and *inhbab*, respectively.** The expression pattern across tissues for **(A)** activin Cb (*inhbcb1* and *inhbcb2*) ohnologs, and **(B)** activin Ab (*inhbab1* and *inhbab2*) ohnologs.

