## Supplementary Information for "Gene function is a driver of signaling pathway evolution following whole genome duplication"

Table S1 – Custom NanoString nCounter Gene Expression Probe Sequences

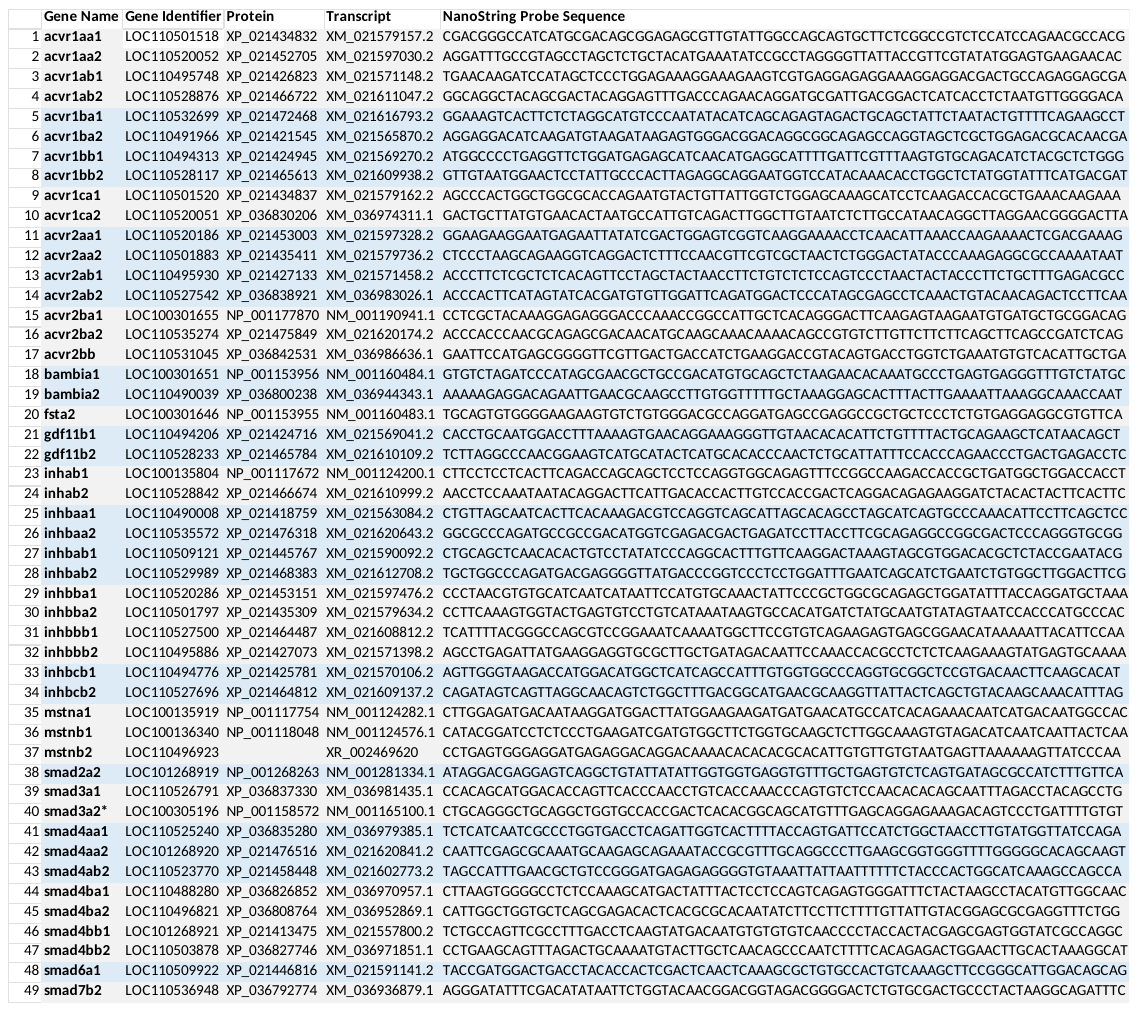

Figure S1

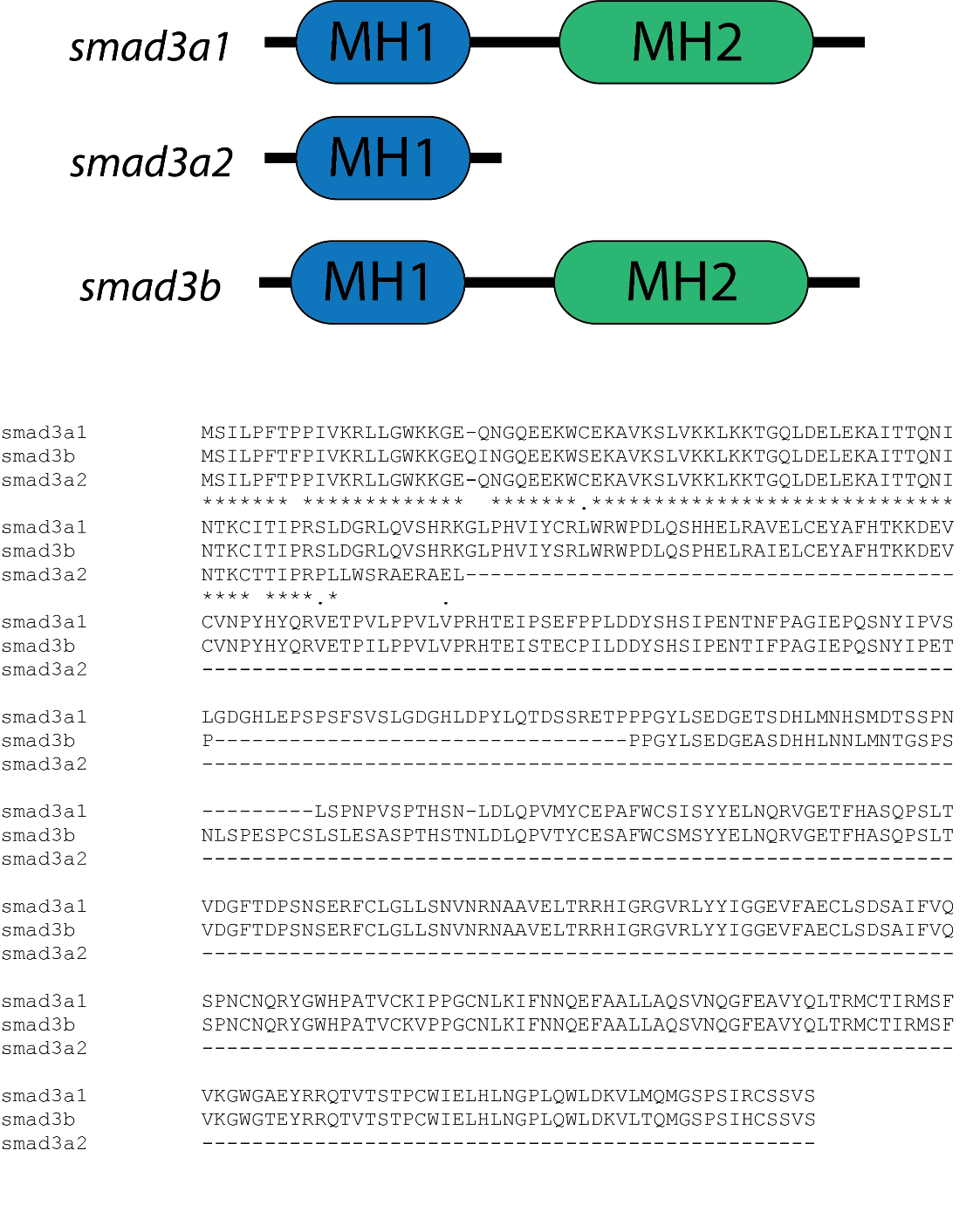

Figure S2

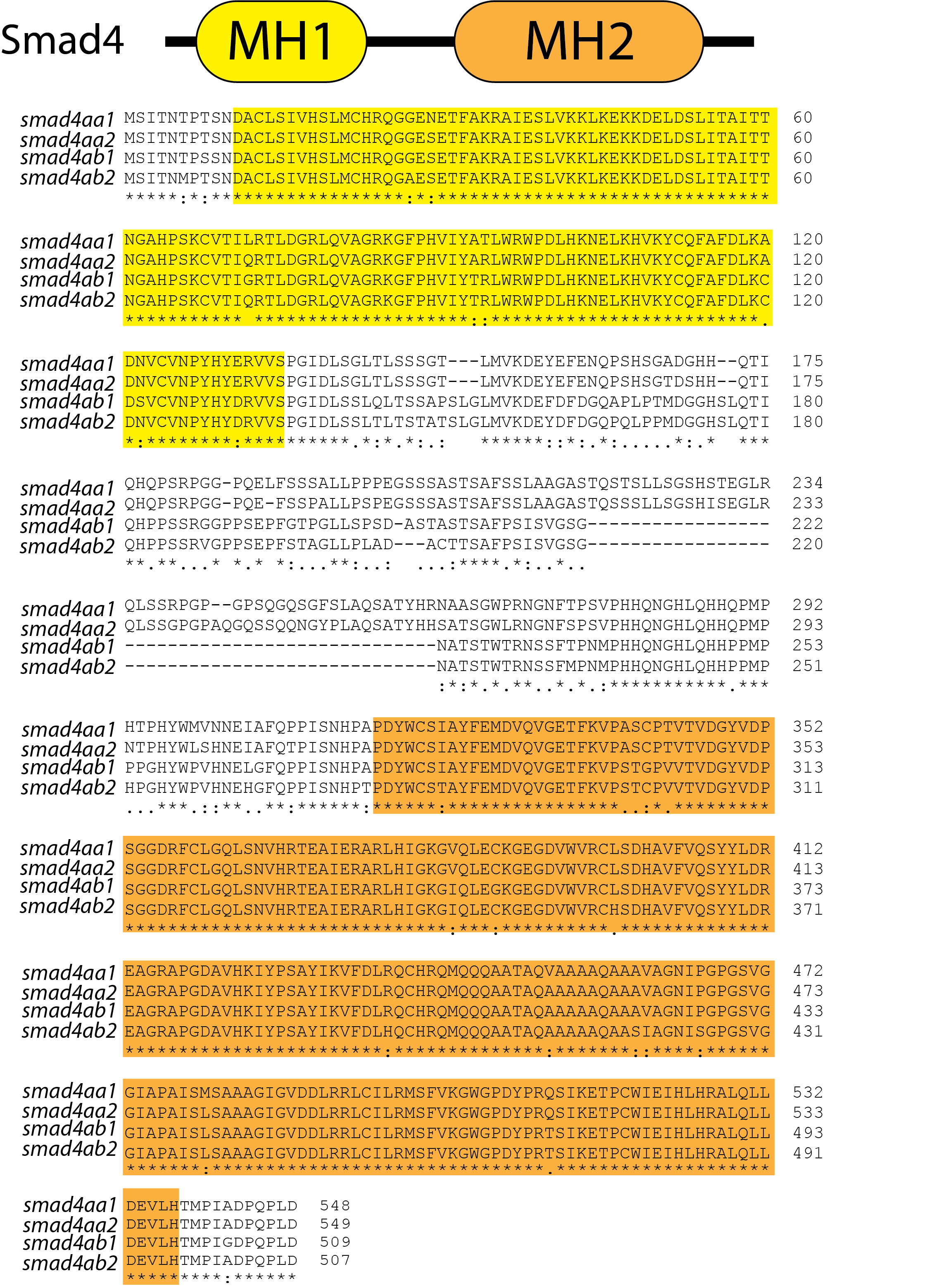

Figure S3

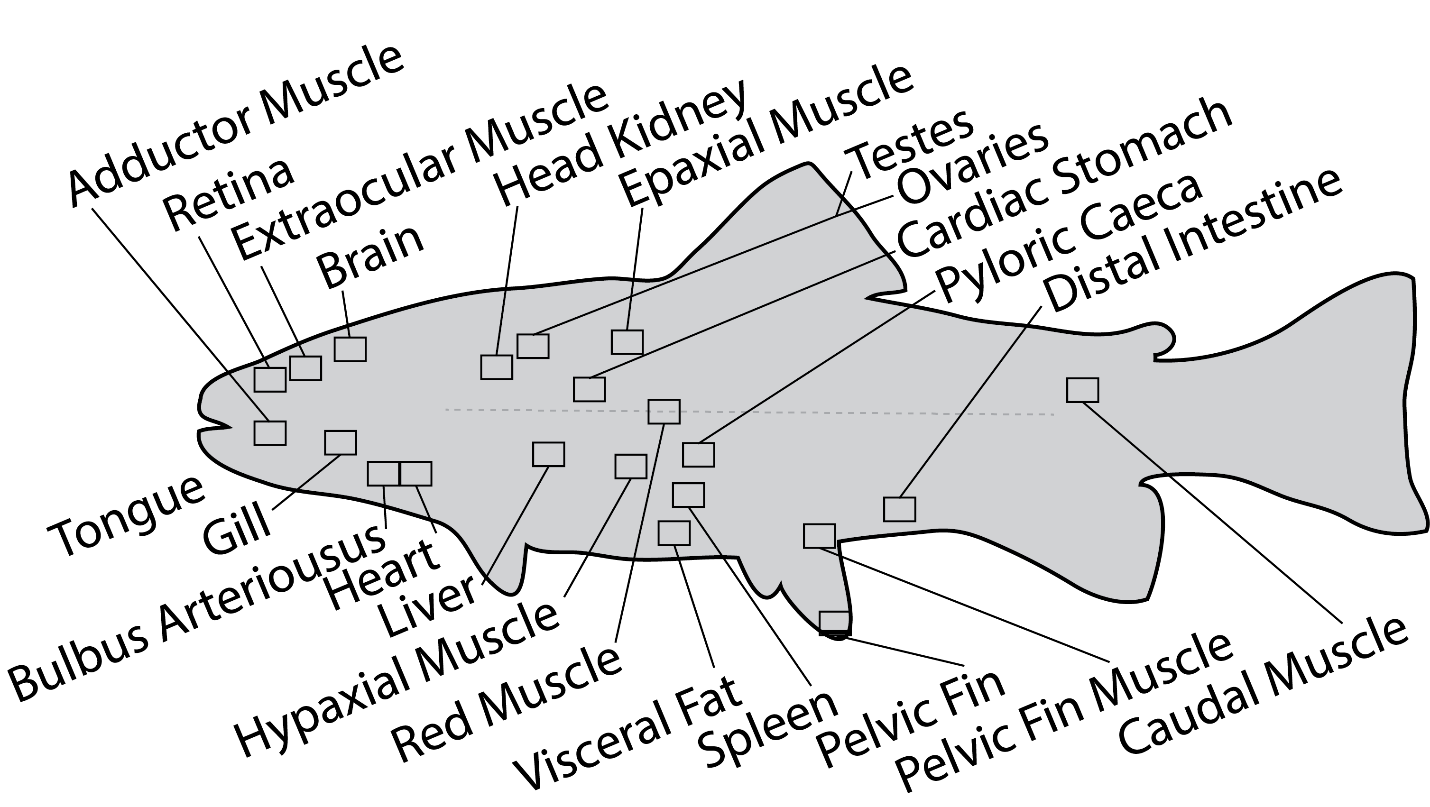

Figure S4

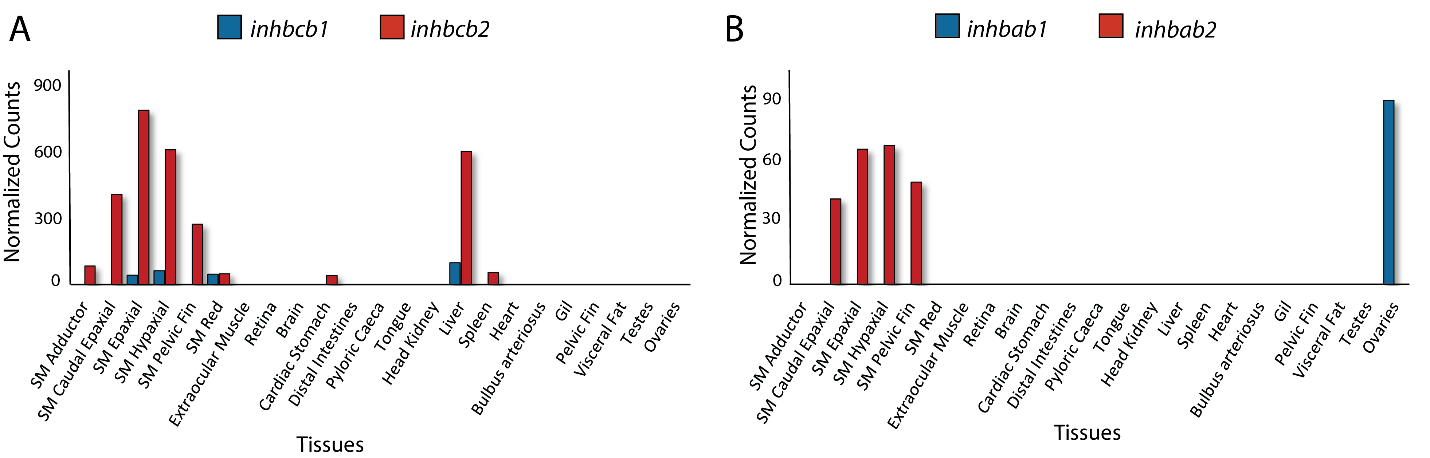
